# Arl3 regulates ODA16-mediated intraflagellar transport in motile cilia biogenesis

**DOI:** 10.1101/2023.04.12.536397

**Authors:** Yameng Huang, Xiaoduo Dong, Stella Y. Sun, Teck-Kwang Lim, Qingsong Lin, Cynthia Y. He

## Abstract

Arl13b and Arl3 are ciliary GTPases implicated in human Joubert Syndrome, affecting ciliary membrane and axoneme organization. Although the mechanism of Arl13b as a guanine nucleotide exchange factor (GEF) of Arl3 and the function of Arl13b and Arl3 in ciliary membrane protein transport are well established, their role in axoneme biogenesis is unclear. In *Trypanosoma brucei*, TbArl13 acts as a GEF for two distinct TbArl3 proteins, TbArl3A and TbArl3C. Here, we identified the *T. brucei* homolog of ODA16, a cargo adapter facilitating intraflagellar transport (IFT) of motile ciliary components, as an effector of both TbArl3A and TbArl3C. Depletion of TbArl3 GTPases stabilized TbODA16 interaction with IFT, while active TbArl3 variants displaced TbODA16 from IFT, demonstrating a mechanism of TbArl3 in motile ciliary cargo transport.

**One-sentence summary:** Arl3 acts as a displacement factor and releases ODA16 from IFT

## Introduction

Cilia and flagella are eukaryotic organelles with motility and sensory functions. Defects in cilia structure and function can result in a large spectrum of diseases known as ciliopathies (*1*). In comparison to primary cilia, motile cilia have additional axonemal structures such as the dynein arms and central pair (CP) microtubules that are required for ciliary beating (*2*). Multiple protein trafficking pathways are involved in cilia biogenesis, maintenance, and function. The best characterized is the intraflagellar transport (IFT) pathway, where the multi-subunit, mega-Dalton IFT complex mediates bidirectional protein trafficking into and out of the ciliary compartment (*3, 4*). Multiple cargo adapters function together with the IFT, facilitating selective transport of a wide range of ciliary cargos (*5*).

Arl13b and Arl3 are cilia-associated, Arf/Arl family GTPases that can be traced back to the last eukaryotic common ancestor (LECA) (*6*). Both small GTPases are implicated in human Joubert syndrome (*7, 8*), and a variety of ciliary functions including ciliogenesis and signaling. Mutations in Arl13b and Arl3 lead to changes in axonemal organization and ciliary membrane protein composition (*9-12*). The seminal discoveries of Arl3 functions in releasing lipidated cargos from carrier proteins Unc119 or PDE6δ (*13, 14*), and that Arl13b acts as a guanine exchange factor (GEF) for Arl3 (*15*) have nicely explained the mechanisms of Arl13b and Arl3 in the transport of selected ciliary membrane proteins involved in signaling (*16*). More recent studies in *Chlamydomonas reinhardtii* have also found a role for Arl13b and Arl3 in BBSome-mediated IFT of ciliary membrane proteins (*17, 18*). The mechanism of Arl13b and Arl3 in axoneme biogenesis, however, is still poorly understood.

Previously, we have identified a single Arl13b ortholog, TbArl13 in the flagellum of *Trypanosoma brucei*, an evolutionarily divergent protozoan parasite that causes Trypanosomiasis (*19, 20*). Two distinct Arl3 homologs, TbArl3A and TbArl3C, interact with TbArl13. Both TbArl3A and TbArl3C can be GTP-loaded by TbArl13 in guanine exchange reactions and both have flagellar functions (*19*). The lipidated cargo carrier TbUNC119 is a specific effector of TbArl3A and functions in lipidated flagellar protein transport (*21*) (Fig. 1A). Interestingly, while TbArl13 is essential for flagellar biogenesis and cell survival (*19*), depletion of TbUNC119 has no observable effects on either flagellar morphology or cell proliferation (*21, 22*). RNAi depletion of TbArl3A or TbArl3C alone moderately affected flagellar morphology but the cells continued to proliferate at a slower rate (Fig. 1, B to D). Simultaneous RNAi of both TbArl3A and 3C, however, led to cell death after 24 hours (Fig. 1, B to D), recapitulating the growth phenotype previously observed with TbArl13 RNAi (*19*). We thus hypothesized that TbArl13 has additional flagellar functions beyond TbUnc119-mediated lipidated protein transport, possibly via additional TbArl3A and TbArl3C effectors (Fig. 1A).

**Fig. 1.**
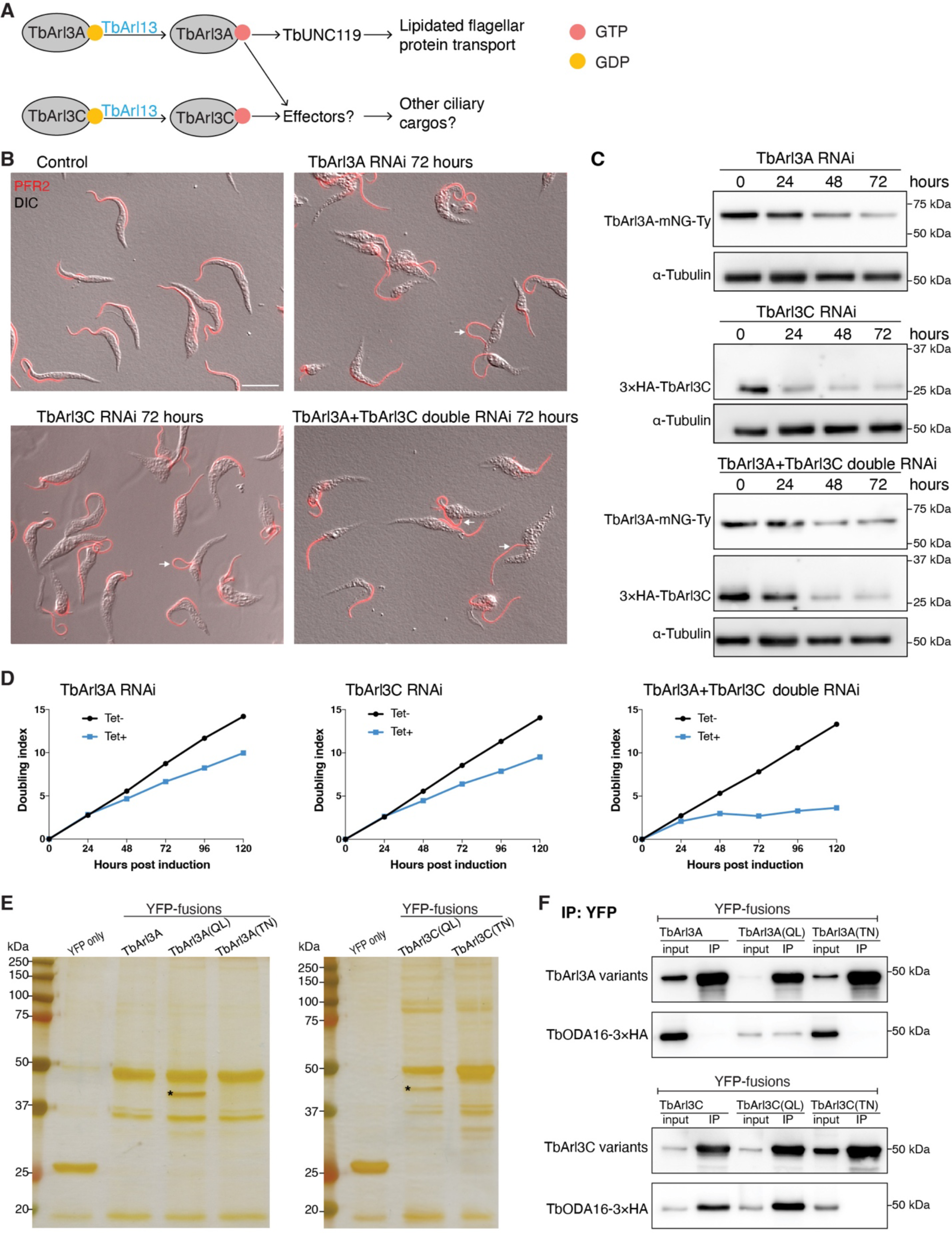
TbODA16 is an effector of both TbArl3A and TbArl3C. (**A**) Overview of the TbArl13-TbArl3 pathway in *T. brucei.* **(B)** *T. brucei* cells were stably transfected with tetracycline (Tet)-inducible RNAi of TbArl3A and TbArl3C, individually or together. In control cells, the flagellum is laterally attached to the cell body. In all RNAi cells, morphology of the flagellum and its attachment to the cell body were affected. Regions of flagellar detachment from the cell body were marked by arrows. Scale bar: 10 μm. (**C**) Efficient depletion of TbArl3A and/or TbArl3C in the RNAi cells was confirmed by immunoblots of endogenously tagged TbArl3A-mNeonGreen (mNG)-Ty and 3×HA-TbArl3C with anti-Ty and anti-HA antibodies. α-tubulin was used as loading control. (**D**) Growth assays of TbArl3A RNAi, TbArl3C RNAi and dual RNAi cells, in the absence or presence of tetracycline for RNAi induction. (**E**) Silver staining of proteins co-immunoprecipitated with TbArl3 variants led to the identification of TbODA16 (marked by *) as a novel effector for TbArl3A and TbArl3C. (**F**) Co-IP experiments showing TbODA16 interaction with TbArl3A and TbArl3C in a GTP-dependent manner.

## Results

### TbODA16 is an effector of TbArl3A and TbArl3C

To search for potential effectors, we expressed QL- and TN-mutants that correspond to constitutively active and inactive forms of TbArl3A and TbArl3C as YFP-fusions using a cumate-inducible system in *T. brucei* (*23*), and performed immunoprecipitation using GFP-Trap. One distinct band (asterisks, Fig. 1E) co-precipitated with the active TbArl3A(Q70L) and TbArl3C(Q77L), but not the inactive TbArl3A(T30N) or TbArl3C(T28N). Mass spectrometry analyses identified this band to be the protein product of Tb927.8.4210 (http://tritrypdb.org). Further bioinformatic analyses showed that Tb927.8.4210 encodes a WD repeat-containing protein that is homologous to ODA16 (Outer Row Dynein Assembly Protein 16 Homolog), which is also known as DAW1 (Dynein assembly factor with WDR repeat domains 1) or WDR69 (WD Repeat-Containing Protein 69). The amino acid sequence encoded by Tb927.8.4210 shares 60% identity with human DAW1 and 65% identity with *C. reinhardtii* ODA16 (Fig. S1). We have therefore renamed Tb927.8.4210 to TbODA16 to reflect this homology. The GTP-dependent interactions between TbArl3 GTPases and TbODA16 were verified by co-immunoprecipitation (co-IP) analyses (Fig. 1F). Interestingly, wild type TbArl3C but not TbArl3A co-precipitated efficiently with TbODA16. It is possible that TbArl3C is more abundantly present in the GTP-bound form in *T. brucei*, as TbArl3C exhibits greater intrinsic GDP-dissociation and GTP-binding activities than TbArl3A (*19*).

### TbODA16 is required for axonemal assembly

ODA16 is best characterized in *C. reinhardtii* (*24-26*), where it is proposed to act as an IFT cargo adapter facilitating the ciliary transport of outer dynein arm (ODA) complex (*27*). The IFT transport of ODA intermediate chain 2 (IC2) was reduced in a *C. reinhardtii* ODA16 mutant (*24, 28*). In *T. brucei*, TbODA16 fusion to mNeonGreen was enriched at the basal bodies when expressed at the endogenous level (Fig. 2A) (*29*). RNAi silencing of TbODA16 slowed cell proliferation in culture (Fig. 2, B and C) and led to reduced flagellar length (Fig. 2, D and E). To further understand the effects of TbODA16 on flagellar assembly, we processed TbODA16 RNAi cells for transmission electron microscopy (TEM) (Fig. 2F). In trypanosome flagellum, the microtubule axoneme is stably associated with a paraflagellar rod (PFR) complex via microtubule doublets 4 to 7 (*30*), providing a convenient positional reference. The orientation of the CP microtubules is fixed, and they always align approximately with doublets 3 and 8 (*30*). In TbODA16 RNAi cells, however, the orientation of the CP microtubules became highly variable. Apart from the CP microtubules, the overall “9+2” axonemal structure appeared intact in TbODA16 RNAi cells. The ODA complex, a known ODA16 cargo, showed no obvious defects. However, the current analyses did not allow for the detection of more subtle flaws.

**Fig. 2.**
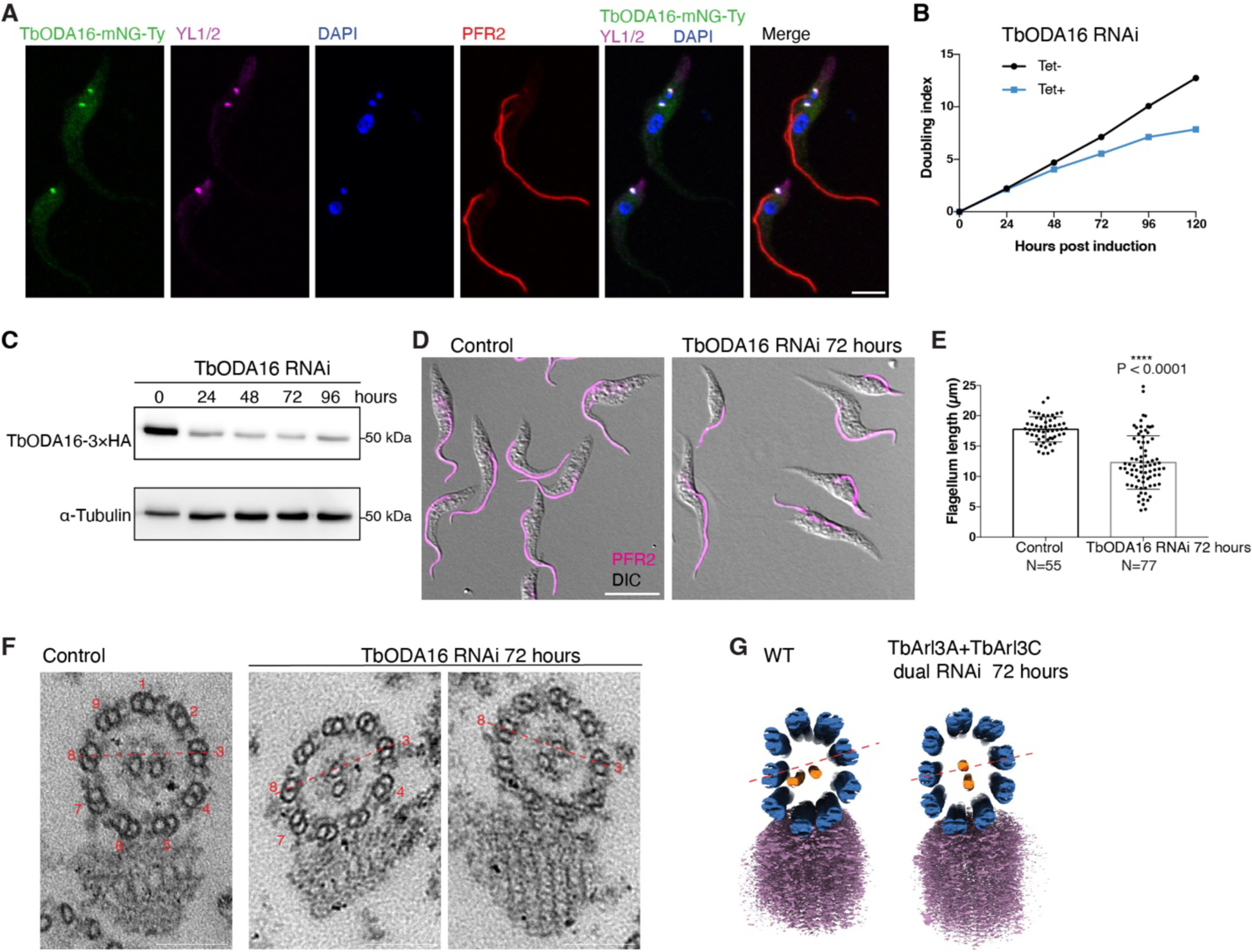
TbODA16 is essential for *T. brucei* cell proliferation and cilia biogenesis. (**A**) Immunofluorescence of cells stably expressing TbODA16 fusion with mNeonGreen from an endogenous allele. The cells were co-stained with anti-YL1/2 for the basal bodies, anti-PFR2 for the flagella and DAPI for the DNA containing nuclei (large ovals) and kinetoplasts (small dots). Scale bar: 5 μm. (**B**) Growth assays of control and TbODA16 RNAi cells. (**C**) Immunoblots showing depletion of endogenously tagged TbODA16-3×HA protein upon induction of TbODA16 RNAi with 10 μg/mL tetracycline. (**D**) Immunofluorescence of control and TbODA16 RNAi cells stained for the flagellum. Scale bar: 10 μm. (**E**) Flagellum length measurement of cells shown in **D**. The results were shown as mean ± SD. P values were calculated by unpaired t test with Welch’s correction. (**F**) Representative TEM images showing altered central pair microtubule position in TbODA16 RNAi cells. Scale bars: 100 nm. (**G**) Representative cryo electron tomography reconstructions showing central pair microtubule (orange) misalignment in TbArl3A/TbArl3C dual RNAi cells. Microtubule doublets and PFR are shown in blue and purple, respectively.

To test if TbODA16 depletion may have changed the molecular composition of the ODA complex, the ODA intermediate chains TbIC1 (also known as TbDNAI1) and TbIC2 (*31*) were tagged with mNeonGreen and expressed endogenously from their native alleles. While TbIC2 localized to the flagellum in both control and TbODA16 RNAi cells without discernible difference (Fig. S2, A and B), TbIC1 intensity along the flagellum was partially reduced in RNAi cells (Fig. S2, C and D). It is unclear why TbODA16 RNAi affected TbIC1 but not TbIC2. The partial reduction of flagellar TbIC1 and unchanged TbIC2 in TbODA16 RNAi cells, however, are consistent with the TEM observation that the ODA complex was generally intact in these cells (Fig. 2F). Similarly in *C. reinhardtii,* zebrafish and human, ODA16 facilitates efficient ODA trafficking, but ODA16 deficiency does not completely inhibit ODA import into the cilia (*24, 32, 33*). Other cargo adapters, for example ODA8 (*34*), may also facilitate flagellar transport of ODA complexes.

The main defect observed in TbODA16 RNAi cells was CP misalignment (Fig. 2F). We found that TbHydin, a known CP component with a role in CP alignment (*35*), was significantly reduced in the flagella upon TbODA16 RNAi (Fig. S2, E and F). The effects of ODA16 mutation on CP alignment have not been reported in other organisms, likely because the CP position is harder to analyze in cilia without a para-axonemal marker such as the PFR; and the CP is observed to rotate relative to the 9 microtubule doublets in some cell types including *C. reinhardtii* (*36*) and *Paramecium* (*37*).

Taken together, our results supported a role of TbODA16 in axoneme assembly, affecting the axonemal localization of TbIC1 and TbHydin, both of which are conserved proteins specifically found in motile cilia. The cellular function of ODA16 in motile cilia biogenesis as previously reported in green algae, zebrafish and mammalian cells is thus also conserved in *T. brucei*.

### ODA16-IFT interaction is regulated by TbArl3A and TbArl3C

Both TbArl3A and TbArl3C interacted with TbODA16 in a GTP-dependent manner (Fig. 1). Cryo-electron tomography analyses of TbArl3A/TbArl3C dual RNAi cells also revealed CP misalignment (Fig. 2G), similar to that observed in TbODA16 RNAi cells. These results suggested a functional connection between TbArl13A/TbArl3C and TbODA16. This connection is highly specific. TbArl2, which is closely related to TbArl3A and TbArl3C (*19*) and affects the microtubule-based cytoskeleton in *T. brucei* (*38*), had no detectable interaction with TbODA16 (Fig. S3A). TbArl3B, another Arl3 homolog in *T. brucei,* does not interact with TbArl13 and does not exhibit any flagella phenotype when overexpressed as a constitutively active mutant (*19*). TbArl3B RNAi also did not have any detectable phenotype on cell growth or flagella morphology (Fig. S3, B to D). As such, in this study we shall focus on TbArl3A and TbArl3C for their potential regulatory functions on TbODA16.

The first hint that TbArl3A and TbArl3C regulate TbODA16 came from localization studies of the latter (Fig. 3A). In control cells, TbODA16 was present at the basal bodies, with some weak signals in the cytoplasm and the ciliary lumen. Upon TbArl3A/TbArl3C dual RNAi, TbODA16 became enriched in the cilia. Ciliary accumulation of TbODA16 was also observed in cells depleted of TbArl3C alone, but not in cells lacking TbArl3A (Fig. S4). As an IFT cargo adapter, ODA16 is expected to eventually dissociate from the IFT and recycle to the ciliary base for another round of cargo transport. Thus, the ciliary accumulation of ODA16 upon TbArl3A/TbArl3C dual RNAi suggested a possible role for TbArl3A and TbArl3C in regulating ODA16 interaction with the IFT train. To investigate this possibility, we performed proximity-based BioID in cells with or without TbArl3A and TbArl3C, using TbODA16-BioID2 as a bait. Interestingly, several IFT subunits including TbIFT46, IFT81 and IFT144 were found more abundantly present in TbArl3A/TbArl3C dual RNAi cells (Fig. 3B). To test if TbODA16-IFT interaction was enhanced in the absence of the TbArl3 GTPases, TbIFT88, TbIFT46, TbIFT20 and TbIFT81 were each tagged endogenously with YFP in cells co-expressing TbODA16-3×HA at the native level. These cells were then induced for TbArl3A/TbArl3C dual RNAi or not, and co-IPs were performed using anti-HA or GFP-Trap (Fig. 4; Fig. S5). In all cases, TbODA16-IFT interaction was stronger in cells depleted of TbArl3A and TbArl3C. Consistent with the above, TbODA16 was mainly present at the basal bodies in control cells; upon depletion of TbArl3A and TbArl3C, TbODA16 became enriched in the cilia, colocalizing with the IFT components (Fig. 4; Fig. S5).

**Fig. 3.**
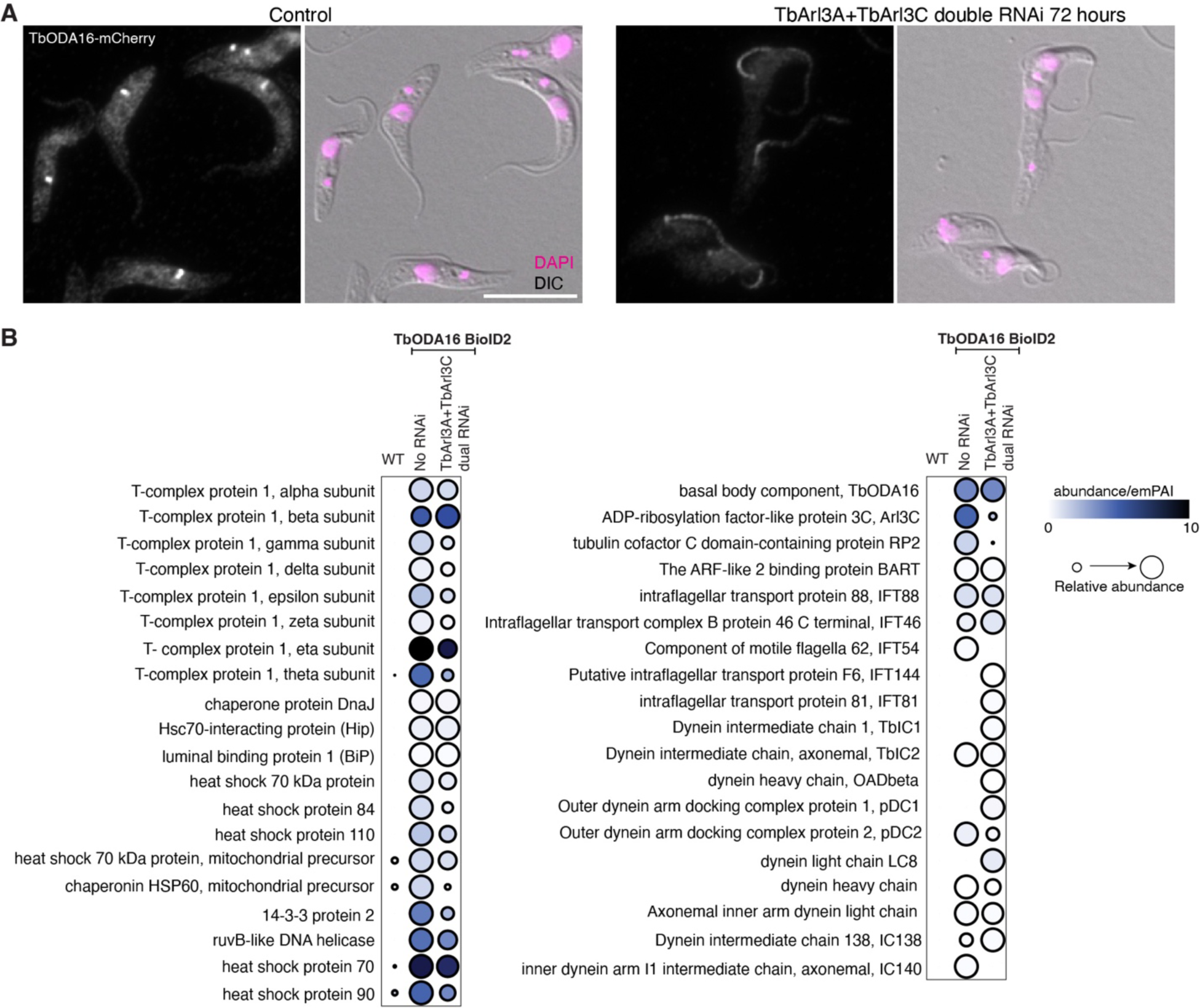
TbArl3 GTPases affect TbODA16 localization and proximity interaction. (**A**) Immunofluorescence of cells stably expressing TbODA16-mCherry, before and after induction for TbArl3A/TbArl3C dual RNAi. Scale bar: 10 μm. (**B**) A comparison of BioID2-based proximity interactomes of TbODA16 before and after TbArl3A/TbArl3C dual RNAi.

**Fig. 4.**
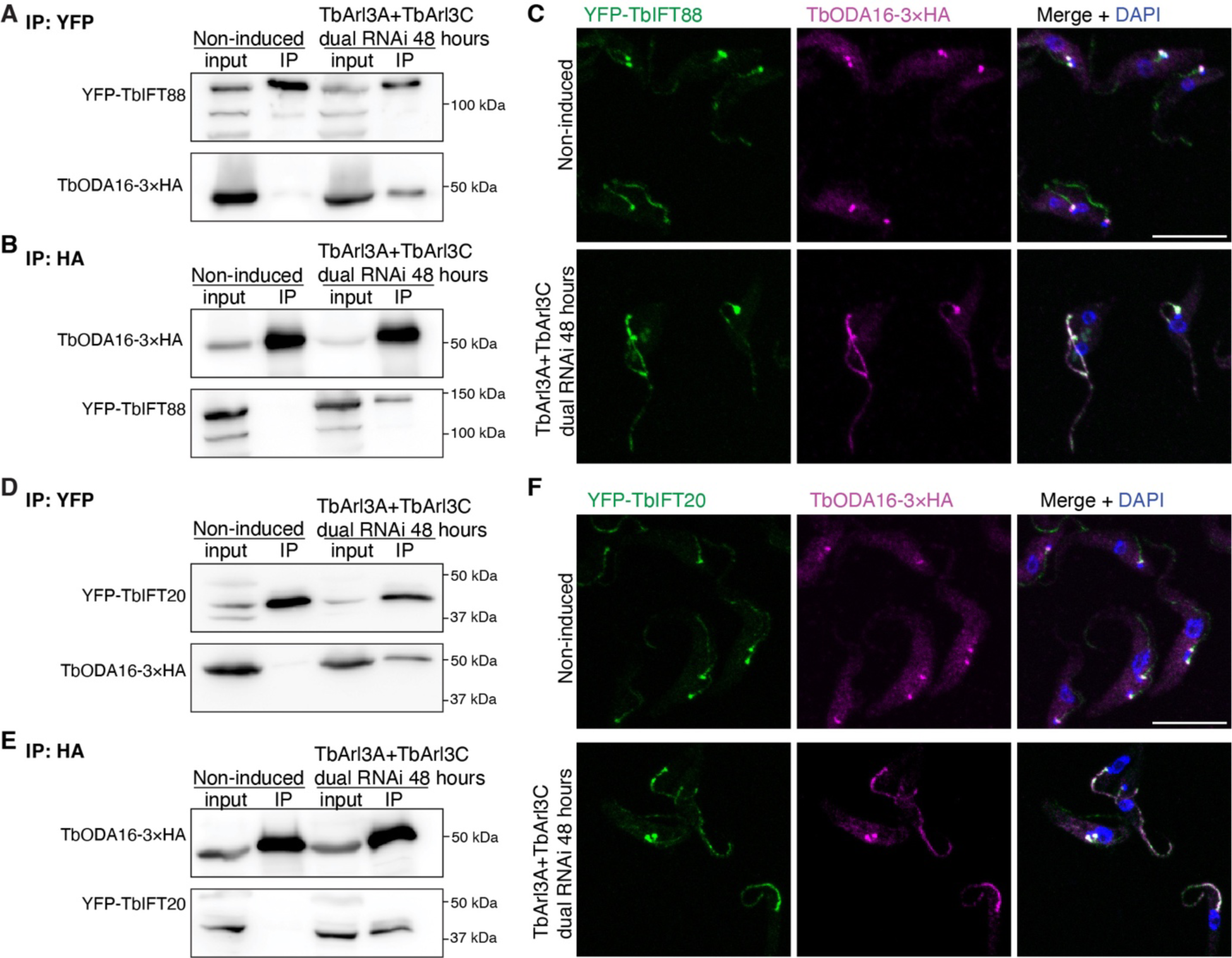
The absence of TbArl3A and TbArl3C stabilizes TbODA16 interaction with IFT. Cells stably expressing HA-tagged TbODA16 and YFP-tagged IFT subunits IFT88 (**A-C**) and IFT20 (**D-F**), respectively, were either un-induced or induced for TbArl3A/TbArl3C dual RNAi. TbODA16-IFT interaction was assessed by co-IP using beads targeting YFP (**A, D**) or HA (**B, E**) tags. Immunofluorescence showing TbODA16 and IFT components in control and TbArl3A/TbArl3C dual RNAi cells (**C, F**). Scale bars: 10 μm.

As both TbArl3A and TbArl3C can be regulated by TbArl13 (*19*), the effects of TbArl13 on TbODA16 were also examined. TbArl13 RNAi led to TbODA16 accumulation in the flagellum, colocalizing with TbIFT88 (Fig. 5A). TbODA16 expression levels were not affected (Fig. 5B). Based on these results, we concluded that TbArl13, TbArl3A and TbArl3C all regulate TbODA16 interaction with the IFT train.

**Fig. 5.**
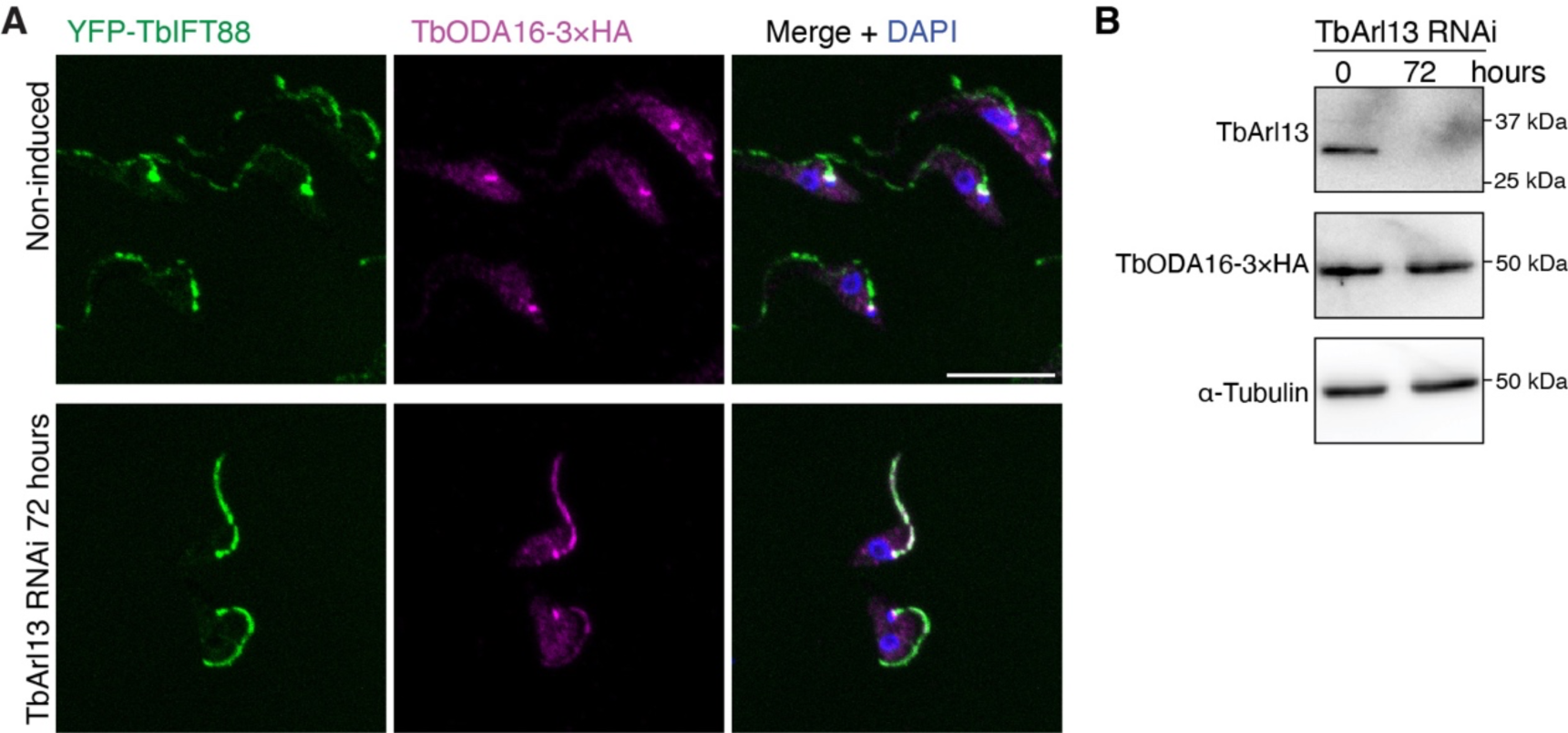
Depletion of TbArl13 leads to TbODA16 accumulation in cilia. (**A**) Representative immunofluorescence images of cells stably expressing HA-tagged TbODA16 and YFP-tagged IFT88, induced for TbArl13 RNAi or not. Scale bar: 10 μm. (**B**) Immunoblots showing depletion of TbArl13 protein, and unchanged expression of TbODA16 upon TbArl13 RNAi.

### Active TbArl3A and TbArl3C can displace TbODA16 from the IFT complex

To test if active TbArl3A and TbArl3C could displace TbODA16 from the IFT complex, the TbArl3A/TbArl3C dual RNAi cell line with stable expression of TbODA16-3×HA and YFP-tagged TbIFT88 was further engineered to include cumate-inducible expression of RNAi-resistant TbArl3 variants. Co-IP was then performed using GFP-Trap and the amount of TbODA16 associated with IFT was evaluated before and after TbArl3 variants were induced (Fig. 6A). In scheme 1, expressions of TbArl3A and TbArl3C variants were induced at the same time of RNAi. TbODA16-IFT interaction was barely detectable in cells expressing TbArl3A and TbArl3C, either wild type or QL forms, demonstrating efficient phenotype rescue to control levels by the RNAi-resistant, active TbArl3 variants. Expression of the inactive TN mutants, however, did not reduce TbODA16-IFT interaction (Fig. 6B). In scheme 2, TbArl3A/TbArl3C dual RNAi was first induced for 30 hours, which stabilized TbODA16-IFT interaction (Fig. 6C). Expression of TbArl3C variants were then induced in the cell system. TbArl3C(QL) rapidly displaced TbODA16 from the IFT within 1 hour of induction. In contrast, TbArl3C(TN) that was expressed at similar levels to TbArl3C(QL) did not affect TbODA16-IFT interaction.

**Fig. 6.**
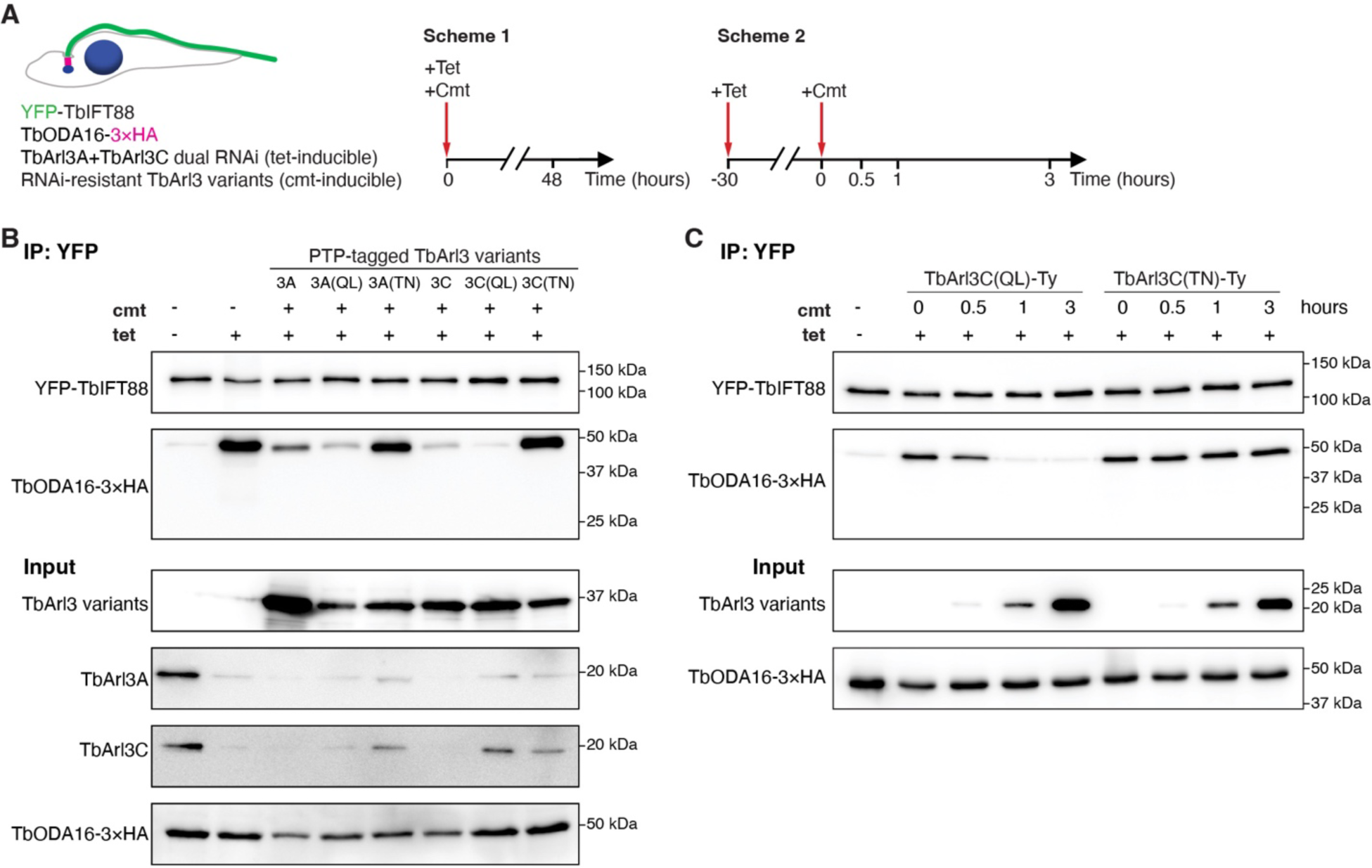
Active TbArl3A and TbArl3C displace TbODA16 from IFT. (**A**) Schematics of the displacement assays. (**B**) Co-IP demonstrating the effects of RNAi-resistant TbArl3 variants on TbODA16-IFT interactions following scheme 1. Tetracycline (Tet) concentration: 10 µg/ml. Cumate (Cmt) concentration: 10 µg/ml. (**C**) Co-IP experiments following scheme 2 demonstrating rapid displacement of TbODA16 from IFT by active TbArl3C. Cmt concentration: 20 µg/ml.

Despite extensive efforts using different tags and different expression systems, purified TbODA16 and individual IFT subunits were highly unstable, making it difficult to test the displacement model *in vitro*. In our TbODA16-BioID analyses, we found TbODA16 to be associated with many chaperone proteins, including all 8 subunits of the T-complex protein Ring Complex (TRiC) (Fig. 3B; Fig. S6A). The interaction between TbODA16 and TRiC was confirmed by co-IP (Fig. S6B). TRiC has been shown to be responsible for the folding of ∼10% of the proteome including tubulin and actin in various eukaryotic organisms. It thus appears that additional cellular factors, particularly chaperones, are required to stabilize TbODA16 and/or its interactions with the IFT complexes and the axonemal cargos.

## Discussion

In this study using *T. brucei* as a motile cilia model, we have demonstrated a role for TbArl13 and TbArl3 GTPases in the regulation of ODA16-mediated IFT, thus affecting ciliary transport of axonemal cargos important for motility. Our findings explain the diverse and essential functions of Arl13b and Arl3 GTPases, as they affect ciliary membrane proteins essential for signaling as well as axonemal components important for motility, via different Arl3 effectors. Our findings also provide a mechanism of ciliary cargo unloading from the IFT, in which active Arl3 functions as displacement factor to release ODA16 from the IFT train (Fig. 7). We speculate that this displacement occurs near the proximal region of the cilia, where TbArl13 is enriched (*19*), allowing released TbODA16 to be immediately recycled to the ciliary base (Fig. 7). IFT cargo unloading is generally thought to occur near the cilia tip, likely when the IFT is remodelled from anterograde to retrograde IFT (*39*). But exceptions have been observed. Live cell imaging of the IFT cargo DRC4 shows that DRC4 can be dissociated from the IFT along the cilia prior to reaching the tip (*40*). The ODA complex also appears to be released from the IFT immediately upon cilia entry, as it does not interact with the IFT in the ciliary compartment (*24, 41*).

**Fig. 7.**
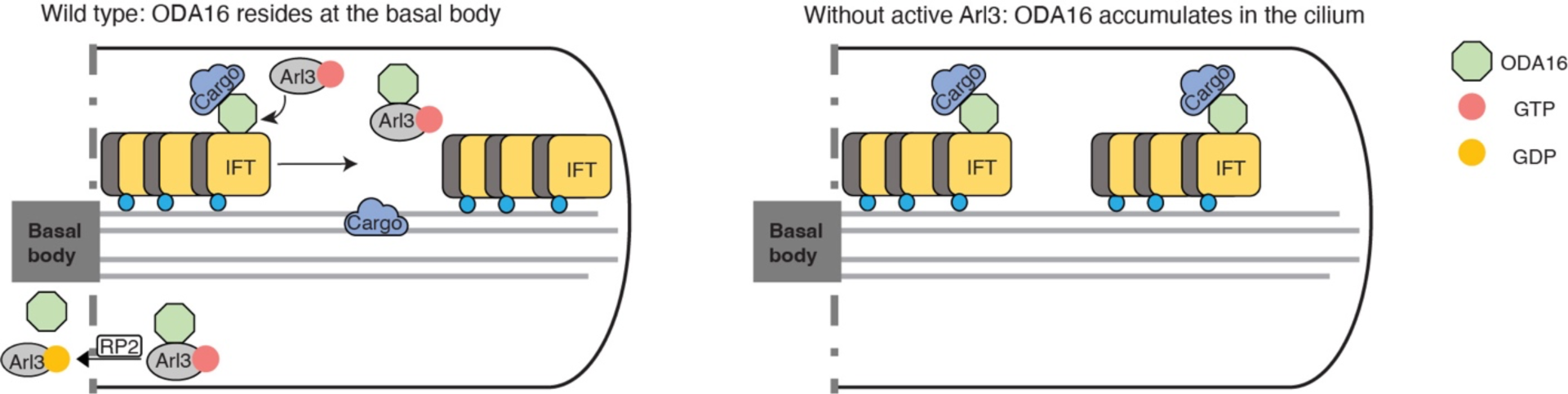
Model for Arl3 in releasing ODA16 from IFT. In wild type cells, ODA16 facilitates ciliary cargo transport by IFT. Upon entry into the cilia, active Arl3 binds to ODA16 and displaces ODA16 from IFT. Hydrolysis of Arl3•GTP by RP2 (*78*), a known Arl3 GTPase-activating protein (GAP) present at the ciliary base (*79*), helps to recycle ODA16 for another round of cargo transport. In cells without active Arl3, ODA16 remains bound to IFT.

Both TbArl3A and TbArl3C, in their respective active forms, were able to regulate ODA16-IFT interactions. However, it appeared that TbArl3C plays a major regulatory role in this regard given its stronger binding to TbODA16 (Fig. 1F) and more pronounced effect on TbODA16 ciliary accumulation upon RNAi (Fig. S4). TbArl3A, in addition to its role in TbUnc119-mediated lipidated protein transport, may partially complement TbArl3C in regulating ODA16-mediated IFT transport. How TbArl3A may act as displacement factor for both lipidated and axonemal cargos is unclear. Previous work demonstrate how Arl3 allosterically regulate prenylated and myristoylated cargo release from PDE6δ and Unc119, through different mechanisms (*13, 14*). Further structural analyses of TbArl3A interaction with TbODA16 will be of particular interest.

ODA16 is the first IFT cargo adapter identified. It aids the IFT transport of the ODA complex (*24, 28*). Direct interaction between ODA16 and ODA components or other ciliary cargos however, has not been demonstrated. In this study, we showed that the flagellar association of two axonemal components, TbIC1 and TbHydin, are reduced in TbODA16 RNAi cells. Depletion of TbIC1 and TbHydin has both been found to cause CP misalignment phenotypes (*30, 35*). These observations thus confirmed the conserved role of ODA16 in motile cilia biogenesis in *T. brucei*. In our TbODA16-BioID analyses, we also found TbODA16 to be associated with several motile ciliary components including TbIC1 and TbIC2 (Fig. 3B). Further examination of BioID candidates in this category will help to identify axonemal cargos transported by TbODA16.

As an IFT cargo adapter, the interaction between ODA16 and IFT have been extensively characterized. In *Chlamydomonas*, ODA16 interacts with the N-terminal 147 amino acids of IFT46 (*24, 25*), and both ODA16 and IFT46 are involved in the flagellar targeting of the ODA complex (*24, 42*). Interestingly, the interaction between DAW1 (human ODA16 homolog) and IFT46 is not conserved in humans (*43*). It is thus possible that the ODA16-IFT interaction is mediated by other IFT components or requires other cellular factors in different organisms. In this study, purified TbODA16 and individual TbIFT components were highly unstable *in vitro*. Therefore we have not been able to test direct interactions between purified TbODA16 with the IFT or the putative axonemal cargos. Our results however, suggest a possibility that TbODA16 works together with other cellular factors, particularly chaperones, in formation and/or stabilization of ODA16 interaction with large protein complexes such as the IFT and the axonemal cargos. ODA16 is recently classified as one of 18 dynein axonemal assembly factors (DNAAFs), which assist the cytoplasmic pre-assembly, maturation and ciliary transport of the axonemal dynein complexes (*44*). The predicted molecular weight of ODA16 homologs from different organisms ranges between 45 to 50 kilodaltons (Fig. S1B). The association of ODA16 with chaperone complexes can also help to explain how a “tiny” cargo adapter could link mega-Dalton complexes such as the ODA and the IFT together.

Arl13b has long been known to affect IFT transport and IFT stabilization in various organisms (*12, 45, 46*), but the mechanisms remain elusive. The current study describes the molecular connection between Arl13b and IFT via a novel Arl3 effector ODA16, and for the first time uncovers the mechanism of Arl13b and Arl3 in motile cilia biogenesis. Notably, a human homolog to Shulin, a *Tetrahymena* DNAAF required for ODA packaging (*47*) is found to interact with Arl3 in mammalian cells containing primary cilia (*48*). The significance of this interaction and whether this interaction also occurs in motile cilia has yet to be determined. Besides Arl13b and Arl3, IFT complex, Hydin and the ODA complex are confirmed human ciliopathy genes (*1*). Recent analyses of ODA16 homologs in mice and humans also demonstrate a link of this gene to primary ciliary dyskinesia, congenital heart diseases and other motile ciliopathies (*33, 49, 50*). The current study thus also shed light on the disease mechanisms of these genes.

## Materials and Methods

### Cell culture

Procyclic form of *Trypanosoma brucei* that proliferates in tsetse fly midgut was used throughout this study. The cells were cultured in Cunningham’s medium (*51*) supplemented with 10% heat-inactivated fetal bovine serum (Hyclone) at 28℃. The cell line DIY (Double-Inducible YTat1.1) was engineered by stable transfection of YTAT1.1 cells (*52*) with a pSmOxNUS vector that enables tetracycline- and cumate-inducible expression (*23*). *T. brucei* strain 427 29-13 (*53*), a tetracycline-inducible cell line, was utilized in TbArl13 RNAi experiments. To select and maintain stable transfectants, the following antibiotic concentrations were applied: 15 µg/ml (for maintenance) or 60 µg/ml (for selection) geneticin, 50 µg/ml hygromycin, 5 µg/ml puromycin, 10 µg/ml blasticidin, 5 µg/ml phleomycin. Unless otherwise stated, 10 µg/ml tetracycline and 10 µg/ml cumate were used to induce gene expressions..

### Transfection

For stable transfection of *T. brucei*, ∼5×10^7^ cells were washed once with 5 ml cytomix (2 mM EGTA pH 8.0, 120 mM KCl, 0.15 mM CaCl_2_, 10 mM K_2_HPO_4_, 5 mM MgCl_2_, 25 mM HEPES, pH 7.6, modified from (*54*)). The cells were then resuspended with 0.5 ml cytomix and combined with 15 µg of linearized DNA. The mixture was transferred to a 0.4 cm electroporation cuvette (BioRad), and electroporated twice at 1500 V, 25 µF, ∞ Ω with 10-second interval using a BioRad Gene Pulser. Clones of stable transfectants were obtained by serial dilution and antibiotic selection for 12-14 days.

### Plasmids

For cumate-inducible expression, the coding sequence of protein-of-interest fused to specified reporter was cloned in pDEX-CuO vectors (*23*). BioID2 was amplified from MCS-BioID2-HA (*55*) (Addgene plasmid # 74224). The PTP tag (ProtC-TEV-ProtA) consists of protein C fusion to protein A, separated by a TEV cleavage site. All constructs listed below are named after the gene, the reporter that each construct contains, and the selectable antibiotics used to generate stable transfectants: TbArl3A-YFP (geneticin), TbArl3A(Q70L)-YFP (geneticin), TbArl3A(T30N)-YFP (geneticin), TbArl3C-YFP (geneticin), TbArl3C(Q77L)-YFP (geneticin), TbArl3C(T28N)-YFP (geneticin), TbODA16-5×GS-3×HA-BioID2 (geneticin), TbODA16-5×GS-YFP (geneticin), GFP-Ty (blasticidin),TbArl3AiR-PTP (hygromycin), TbArl3AiR(Q70L)-PTP (hygromycin), TbArl3AiR(T30N)-PTP (hygromycin), TbArl3CiR-PTP (hygromycin), TbArl3CiR(Q77L)-PTP (hygromycin), TbArl3CiR (T28N)-PTP (hygromycin), TbArl3CiR(Q77L)-Ty (hygromycin), TbArl3CiR (T28N)-Ty (hygromycin), PTP-TbArl2 (blasticidin), PTP-TbArl2(Q70L) (blasticidin), PTP-TbArl2 (T31N) (blasticidin), TbArl3A-PTP (geneticin), TbArl3A(Q70L)-PTP (geneticin), TbArl3A(T30N)-PTP (geneticin), TbArl3C-PTP (geneticin), TbArl3C(Q77L)-PTP (geneticin), and TbArl3C(T28N)-PTP (geneticin). For cumate-inducible expression of RNAi-resistant (iR) TbArl3 variants, RNAi-resistant TbArl3A and TbArl3C sequences were designed by Synonymous Mutation Generator (jong2.pythonanywhere.com) (*56*). Synthetic TbTCP-1-zeta (aa376-544), TbArl3AiR and TbArl3CiR sequences are aligned with wild-type (WT) sequences through M-Coffee (*57*) and shown below.

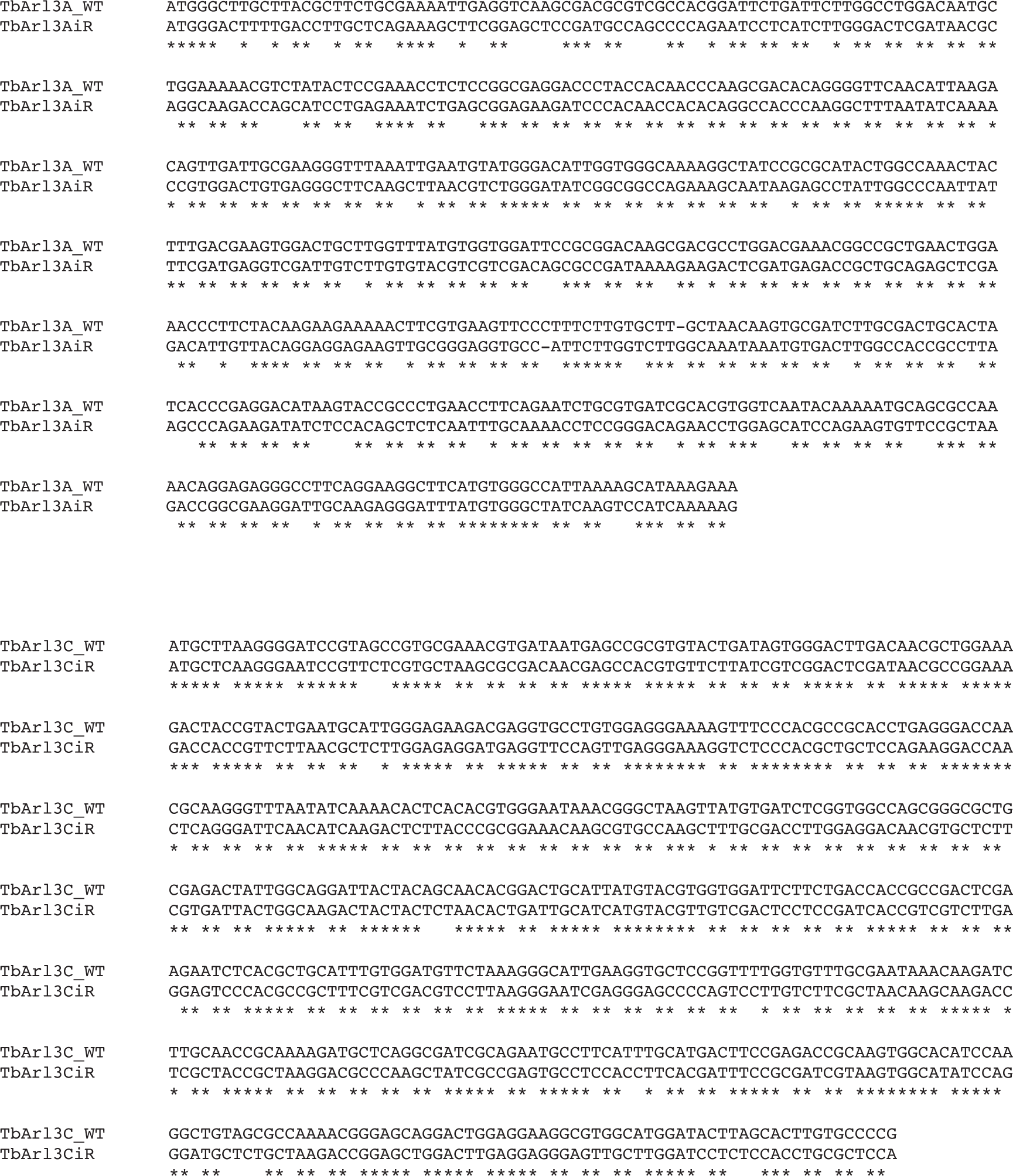

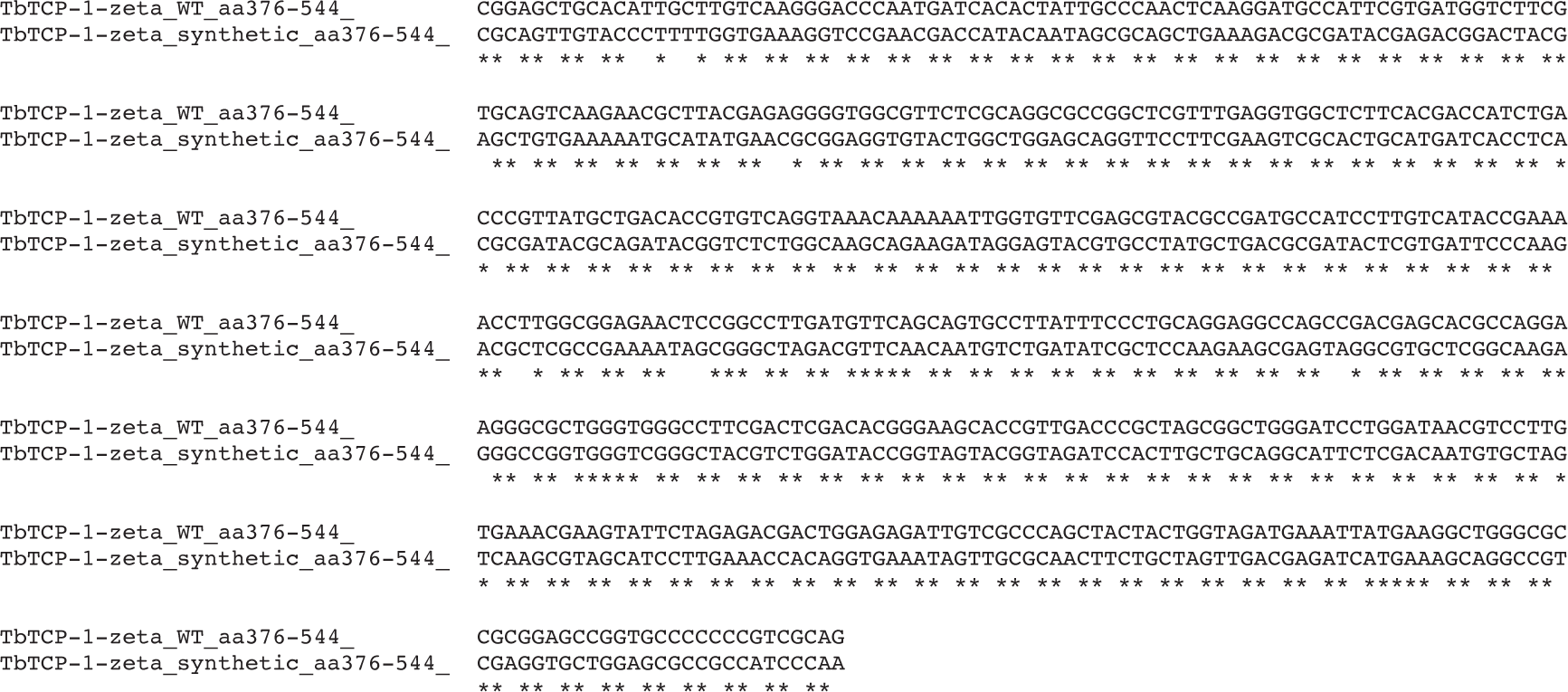

For tetracycline-inducible RNA interference, RNAi targeted sequences (shown in the brackets) were chosen from the RNAit server (https://dag.compbio.dundee.ac.uk/RNAit/) (*58*). The target sequence was then cloned into the p2T7 vector with phleomycin resistance (*59*). The RNAi constructs used in this study are: TbArl3A(nt56-465), TbArl3C(nt38-458), TbArl3C(nt38-458)-TbArl3A(nt56-465) for TbArl3A/TbArl3C dual RNAi, TbArl3B(nt139-547), TbODA16(nt536-1126), TbArl13(nt160-593) (*19*).

For endogenous tagging, desired amplicons were PCR amplified using the pPOT vectors as templates (*60*). Using pPOTv6 and v7 vectors (kind gifts from Keith Gull, Samuel Dean and Jack Sunter) as templates, the following new templates were constructed: pPOTv7-g418-mCherry-g418, pPOTv7-g418-YFP-g418, pPOTv7-blast-3×HA-blast, pPOTv7-hygro-3×HA-hygro, pPOTv7-puro-3×HA-puro, pPOTv6-blast-3Ty::mNG::3Ty-blast, pPOTv6-hygro-3Ty::mNG::3Ty-hygro. TbTCP-1-zeta(aa1-375)::Strep-Tag II::CBP::HA::TbTCP-1-zeta (synthetic aa376-544)-hygro-3’UTR (nt1-861) was constructed by overlapping PCR for endogenously tagging of TbTCP-1-zeta-HA. The amplicons for TbArl3A-mNG-Ty (blasticidin) and TbArl3B-mNG-Ty (blasticidin) contain ∼500 bp of homologous regions. For constitutive overexpression, pXS2-YFP (*61*) was used.

### Cell growth assays

Growth assays were initiated with a freshly diluted culture containing 1×10^6^ cells/ml *T. brucei* cells. Cell density was monitored using a hemocytometer every 24 hours. The cell culture was then diluted with fresh medium to 1×10^6^ cells/ml to maintain the cells in exponential growth phase. Doubling index was calculated as log_2_ (N_t_×D_f_/N_0_), where N_t_ is the cell density at a given time point, D_f_ is the accumulative dilution factor and N_0_ is the cell density at t=0.

### Primary antibodies for immunoblots

The following antibodies were commercially available: anti-HA (mouse, Santa Cruz, sc-7392, 1:500), anti-HA (mouse, BioLegend, 901501, 1:2000), anti-Protein C (mouse, Genscript, A01774, 1:1000), anti-α-tubulin (mouse, Santa cruz, sc-23948, 1:5000), anti-mCherry (rabbit, Invitrogen, PA5-34974,1:3000), Streptavidin-HRP (peroxidase-conjugated streptavidin biotin-binding protein, Thermo Scientific, 21130, 1:10000).

Polyclonal anti-TbArl13 (rabbit, 1:1000) was described previously (*19*). Monoclonal anti-Ty (mouse, 1:500) was a kind gift from Philippe Bastin (*62*). His-PFR2, His-YFP, GST-TbArl3A(Q70L), GST-TbArl3C(Q77L) and MBP-TbODA16 proteins were expressed and purified from *E. Coli* (BL21). Purified proteins were used for customized antibody production by Abnova. Affinity purified antibodies were validated by immunoblots. Listed below are the immunoblot details of the customized antibodies: anti-YFP (rabbit, 1:1000), anti-TbArl3A (rabbit, 1:250), anti-TbArl3C (rabbit, 1:250), anti-TbODA16 (rat, 1:250).

### Primary antibodies for immunofluorescence assays

Anti-PFR2 (rabbit, customized/Abnova, 1:4000), Anti-RFP (rabbit, Rockland, 600-401-379, 1:500), YL1/2 (anti-Tyrosinated α-tubulin, rat, Santa Cruz, sc-53029, 1:2000), anti-HA (mouse, Santa Cruz, sc-7392, 1:250).

### Light microscopy

*T. brucei* cells were washed twice with phosphate-buffered saline (PBS, pH 7.4) before adhering to coverslips by centrifugation (2000g, 1 min). Cells were fixed in 4% paraformaldehyde (PFA) for 10 min, permeabilized with 0.25% Triton X-100 in PBS for 5 min at room temperature, and blocked with 3% BSA (bovine serum albumin) in PBS for 30 min. The cells were then incubated with primary antibodies in the blocking buffer for 1 hour. After three washes with PBS, cells were incubated with Alexa Fluor conjugated secondary antibodies (Invitrogen) and DAPI for 1 hour. Cells were washed twice with PBS and once with distilled water prior to mounting to glass slides. To label the paraflagellar rod with anti-PFR2, cells were fixed and permeabilized with cold methanol at −20°C for 5 min. Images were acquired either by a Zeiss Axio Observer Z1 fluorescence microscope with a 63×/1.4 objective or a FLUOVIEW FV3000 confocal microscope equipped with a U Plan Super Apochromat 60×/1.35 Objective.

### Immunoblot analyses

Samples lysed in Laemmli sample buffer were boiled at 100℃ for 5 min. Proteins were resolved on SDS-PAGE and transferred onto PVDF membranes. Blots were firstly blocked with 3% BSA or non-fat milk in TBST and incubated with the indicated primary antibodies followed by appropriate HRP-conjugated secondary antibodies. Chemiluminescent signal was detected and imaged using ImageQuant LAS 4000 mini (GE Healthcare). Membrane stripping, if required, was done in 0.1 M NaOH for 1 hour.

### Immunoprecipitation and silver staining

3-6×10^8^ *T. brucei* cells were harvested and washed twice with PBS before cell lysis for 15 min on ice in 1 ml lysis buffer (20 mM Tris-HCl pH 7.5, 150 mM NaCl, 1 mM EDTA, 0.5% NP-40) supplemented with 2× protease inhibitor cocktail (PI). Cleared cell lysates were obtained by centrifugation at 17000g for 15 min at 4°C. Supernatants were supplemented with 0.5 ml binding buffer (10 mM Tris-HCl pH 7.5, 150 mM NaCl) and then incubated with GFP-nAb Magnetic Agarose beads also known as GFP-Trap (Allele Biotech or Chromotek), EZview Red Anti-HA Affinity Gel (Millipore) or IgG Sepharose 6 Fast Flow beads (Cytiva) for 90 min at 4°C with rotation. The beads were washed three times in binding buffer containing 0.5× PI, eluted in Laemmli sample buffer and fractionated by SDS-PAGE for further analyses by immunoblots or silver staining. Silver staining was performed according to Blum silver staining protocol (*63*). Protein bands of interest were excised for LC-MS/MS by the Mass Spectrometer Facility at Nanyang Technological University.

### Proximity-dependent biotin identification (BioID)

An optimized 5×GS linker was inserted between TbODA16 and 3×HA-BioID2 reporter to ensure correct cellular localization of the fusion protein. TbArl3A/TbArl3C dual RNAi was induced with 10 µg/ml tetracycline for 24 hours prior to the induction of TbODA16-5×GS-3×HA-BioID2 expression with 10 µg/ml cumate for an additional 24 hours. 50 µM biotin was then added to WT cells, TbODA16-5×GS-3×HA-BioID2 in cells with or without TbArl3A/TbArl3C dual RNAi for 24 hours. For each sample, 4×10^9^ cells were harvested and washed extensively with PBS to remove excess biotin. Cells were then lysed with 2 ml BioID lysis buffer (1% SDS, 500 mM NaCl, 5 mM EDTA, 1 mM dithiothreitol, 50 mM Tris-HCl, pH 7.4) supplemented with 2× PI for 15 min at room temperature. 8 ml 1% NP-40 in PEM buffer with 2× PI was added to the mixture for another 15 min. After centrifugation (16000g, 10 min, 16°C), the supernatant was incubated with 300 μl Dynabeads M-280 Streptavidin (Invitrogen) for 4 hours at 4°C with rotation. Beads were washed once with PBS containing 0.5% SDS, twice with PBS containing 1% NP40 and thrice with PBS. After washing, beads were subjected to disulfide reduction in 200 µl triethylammonium (500 mM, pH 8.5) containing 4 mM Tris (2-carboxyethyl) phosphine (TCEP) for 1 hour at 65°C with gentle agitation, followed by alkylation with addition of 4 μl methyl methanethiosulfonate (MMTS) at room temperature for 15 min. Trypsin was added to 12.5 ng/μl for overnight digestion at 37°C. After trypsin digestion, the peptide solution was separated from beads, desalted and analysed by LC-MS/MS using a TripleTOF 5600 system. Proteomics data were analysed on the ProteinPilot software 5.0 with 1% false discovery rate and searched against Uniprot *T. brucei* proteome and common Repository of Adventitious Proteins (cRAP). Exponentially modified protein abundance index (emPAI) (*64*) was calculated to assess protein abundance.

### Transmission electron microscopy

*T. brucei* cells were firstly extracted with 1% NP40 in PEME buffer (100 mM PIPES pH 6.9, 1 mM MgSO_4_, 0.1 mM EDTA, 2 mM EGTA) for 5 min at room temperature. Subsequently, isolated flagella were washed twice with PEME buffer and fixed with 1ml buffered fixative (2.5% glutaraldehyde, 2% paraformaldehyde, 100 mM phosphate buffer pH 7.0). Resin-embedded *T. brucei* samples were prepared according to established protocols (*65*). Embedded samples were subjected to ultra-sectioning by a Leica Ultracut UCT Ultramicrotome to produce sections below 70 nm. After post staining with UranyLess and Reynolds lead citrate (10min each), these sections were imaged by Tecnai T12 (FEI).

Cryo-electron Tomography

*T. brucei* flagella were isolated following published protocols (*66*) with slight modifications. Cells were extracted in 1% NP40 in PEME buffer supplemented with 1× PI and 0.25 mg/ml DNase I for 10 min at room temperature and subsequent 30 min on ice. Extracted flagella were harvested by centrifugation at 16,000g for 10 min, washed twice with PEME buffer and resuspended in the same buffer. Plasma treated Quantifoil EM grids were mounted on a manual plunger, loaded with isolated flagellum suspension mixed with 10 nm or 15 nm gold fiducials (EMS), blotted from the back side using Whatman paper #5, and plunged into ethane pre-cooled to liquid nitrogen temperature. Isolated flagella were imaged using a 300kV Titan Krios electron microscope (Thermo Fisher) equipped with an energy filter (Gatan), a K3 Summit direct electron detector (Gatan) and an objective aperture or a Volta Phase Plate (*67*). High-magnification images were recorded at 42,000× and 33,000× respectively corresponding to a pixel size 2.202 Å or 2.756 Å. Tilt series were recorded using Tomo4 software with bidirectional acquisition schemes (*68*), each from −50° to 50° with 2° increment. Target defocus was set to −4 µm to −8 µm. The total dose was limited to 90-100 e/Å^2^. Upon phase plate alignment and conditioning, tilt series of the flagellum were recorded at 33,000× at pixel size 2.756 Å using Tomo4 software with bidirectional acquisition schemes, each from −50° to 50° with 2° increment. Target defocus was set to −1.0 µm. Every 2 or 3 tilt series, a new spot on the phase plate was selected. The total dose was limited to 110 e/Å^2^. Tilt series alignment and tomogram reconstruction are performed automatically using tomography pipeline described in (*69*). Subcellular features were semi-automatically segmented using EMAN2 and refined manually using Chimera (*70*) or ChimeraX (*71, 72*).

### Bioinformatic analyses

All *T. brucei* gene sequences were retrieved from tritrydb.org. Sequence information and domain prediction were retrieved from UniProt. The schematic diagram of domains was achieved with IBS web server (*73*). Sequences of ODA16 homologues were aligned with MUSCLE (*74*) and visualized with ESPript (*75*). Secondary structures of HsDAW1 (PDB: 5NNZ) were attached to the alignment for reference.

### Image processing and statistical analyses

Dot plots were generated by ProHits-viz (*76*). Fiji software (*77*) with appropriate plugins was used for image processing. Intensity and length measurements were performed on *T. brucei* cells in early cell cycle stages with a single flagellum. Statistical analysis and graphing were carried out on GraphPad. Sample size (N) and applied analysis were included in corresponding Figure legends.

## Acknowledgements

We thank Tang Bor Luen for critical reading of the manuscript, Tong Yan at The Centre for BioImaging Sciences, National University of Singapore for assistance with confocal microscopy, Sze Siu Kwan and Meng Wei at the Mass Spectrometry core facility at Nanyang Technological University for protein identification by LC-MS/MS, and Yang Hao Yuan and Kwok Yee Ther for technical assistance. This work is supported by Tier 2 research grants (MOE2017-T2-2-109 and MOE-T2EP30121-0003) from Singapore Ministry of Education.

## Author contributions

YH, performed all molecular, biochemical and cell biology assays, and proteomics data analyses; XD and SYS, TEM and cryoET analyses; TKL and QL, proteomics analyses of BioID samples; YH and CYH, conceptualization and writing of the manuscript with the help from all authors.

## Competing interests

The authors declare no competing interests.

## Data and materials availability

All data are available in the main text or the supplementary materials. BioID proteomics data will be deposited to the Proteome-Xchange Consortium via PRIDE. The tomograms will be deposited to EMDataBank database.

## Supplementary Figures

**Fig. S1.**
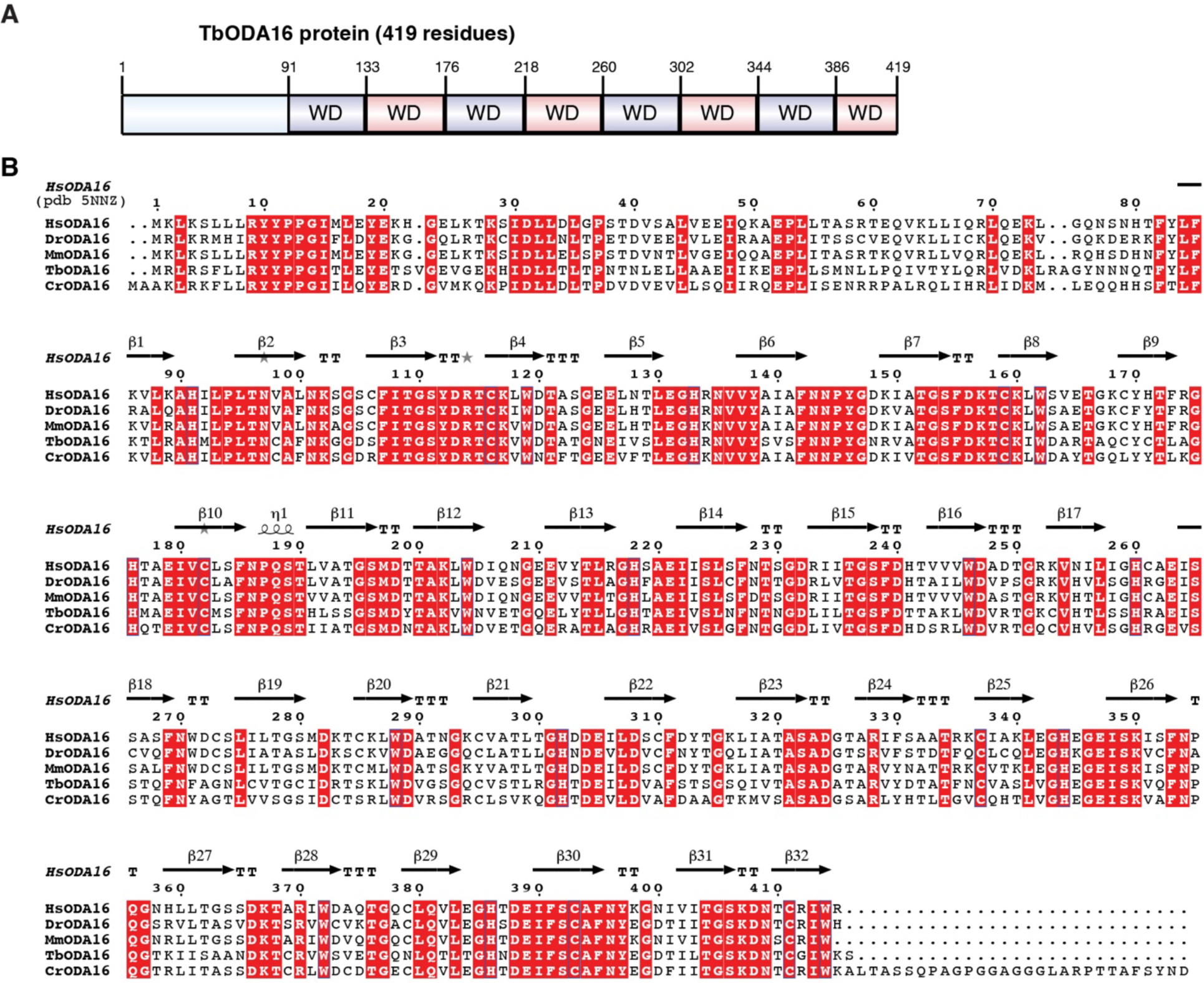
Tb927.8.4210 encodes *T. brucei* homolog to ODA16. (**A**) Tb927.8.4210 encodes a 419-aa polypeptide containing 8 WD repeats. Domain prediction is retrieved from UniProt (Q57W14). (**B**) Multiple sequence alignment of ODA16 homologues from indicated species. Secondary structures of HsODA16 (PDB: 5NNZ) are attached to the alignment as reference. Identical residues are boxed in red. Hs: *Homo sapiens*; Dr: *Danio rerio*; Mm: *Mus musculus*; Tb: *Trypanosoma brucei*; Cr: *Chlamydomonas reinhardtii*.

**Fig. S2.**
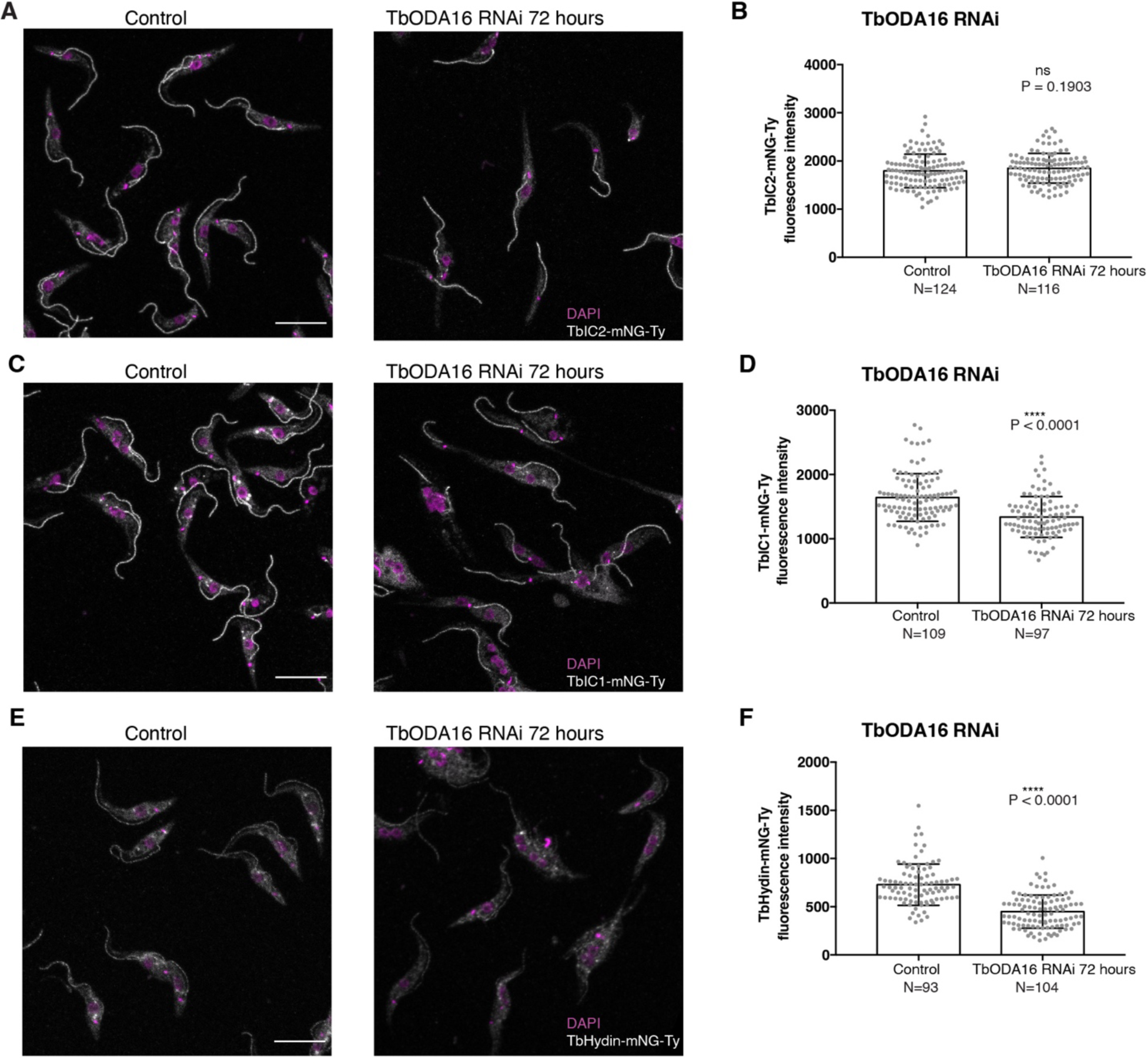
The axonemal association of TbIC1 and TbHydin is reduced upon TbODA16 RNAi. TbODA16 RNAi was induced in cells expressing mNeonGreen fusion to TbIC2 (**A**), TbIC1 (**C**) and TbHydin (**E**). Scale bars: 10 μm. Their flagellar intensity was measured along the distal 1.5 µm and shown in (**B**), (**D**) and (**F**), respectively. The results were shown as mean ± SD. P values were calculated by unpaired t test with Welch’s correction.

**Fig. S3.**
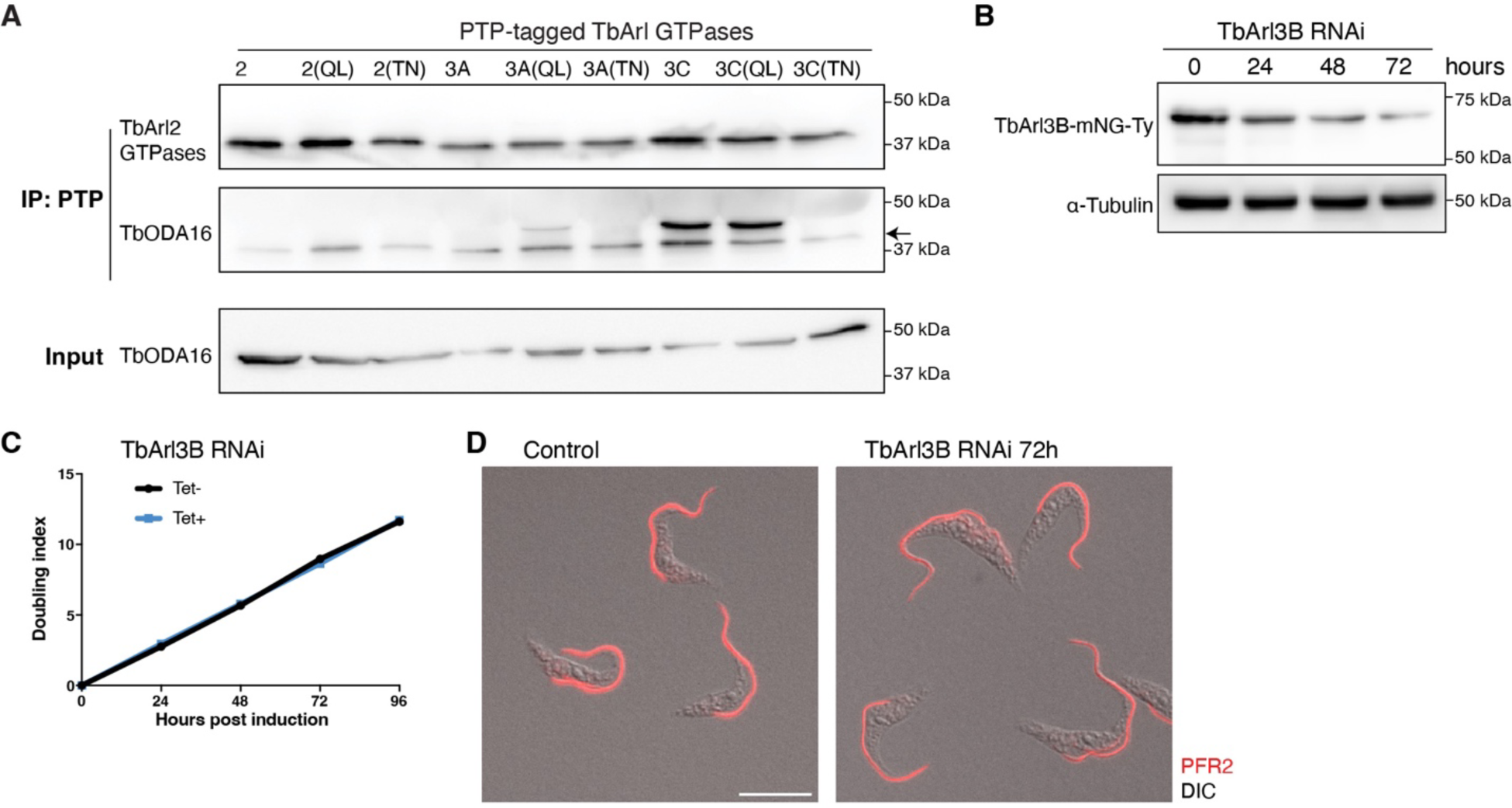
TbArl2 and TbArl3B are not involved in TbODA16 regulation. (**A**) Interaction between TbODA16 and TbArl2 and TbArl3 variants by co-IP assays using IgG Sepharose 6 Fast Flow beads. The anti-TbODA16 antibodies cross-reacted with the PTP tag and therefore also labeled the PTP-tagged TbArl GTPases at ∼40 kDa (arrow). (**B**) Immunoblots showing depletion of TbArl3B by inducible RNAi with 10 μg/mL tetracycline. (**C**) Growth assays of TbArl3B RNAi cells. (**D**) Immunofluorescence of cells non-induced or induced for TbArl3B RNAi for 72 hours. Scale bar: 10 μm.

**Fig. S4.**
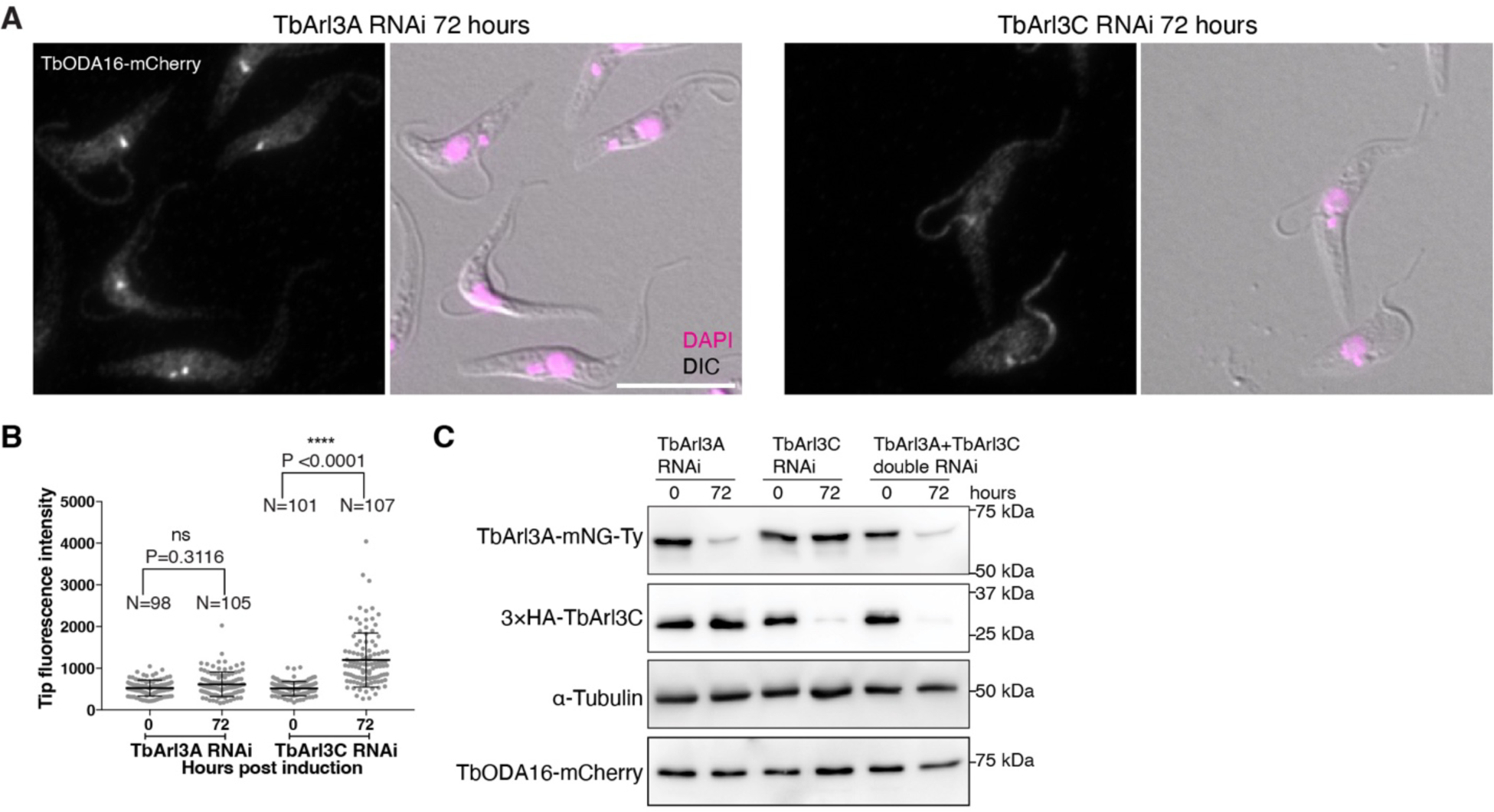
Depletion of TbArl3C but not TbArl3A affects TbODA16 distribution. (**A**) Immunofluorescence of cells with endogenously expressed TbODA16-mCherry, TbArl3A-mNG-Ty and 3×HA-TbArl3C upon induction for TbArl3A RNAi or TbArl3C RNAi for 72 hours. Scale bar: 10 μm. (**B**) Comparison of TbODA16 intensity at the ciliary tip before and after RNAi of specified TbArl3 GTPases. The intensity was measured along the distal 1.5 µm of the flagellum. The results were shown as mean ± SD. P values were obtained from one-way ANOVA with Tukey’s multiple comparisons test. (**C**) Immunoblots confirming efficient and specific TbArl3 depletion in each of the RNAi cell lines shown in this Figure as well as in Fig. 3A.

**Fig. S5.**
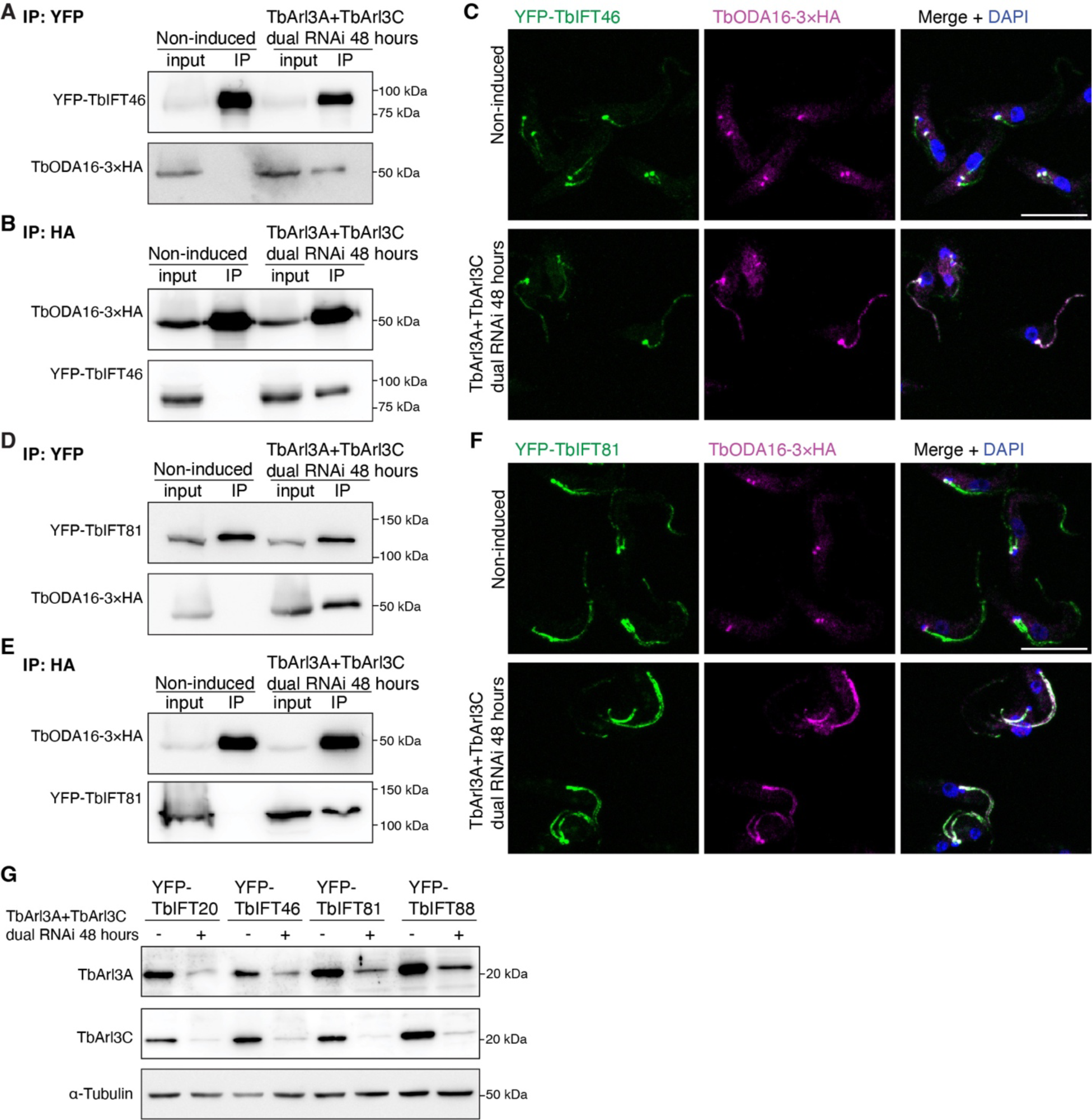
Dual silencing of TbArl3A and TbArl3C stabilizes TbODA16-IFT interaction. (**A-F**) Cells stably expressing HA-tagged TbODA16 and YFP-tagged IFT subunits IFT46 (**A-C**) or IFT81 (**D-F**) were induced for TbArl3A/TbArl3C dual RNAi or not. TbODA16-IFT interaction was assessed by co-IP using GFP-Trap (**A, D**) or anti-HA affinity gel (**B, E**). Immunofluorescence showing TbODA16 in control and TbArl3A/TbArl3C dual RNAi cells (**C, F**). Scale bars: 10 μm. (**G**) Immunoblots confirming the depletion of TbArl3A and TbArl3C in all cell lines used in experiment shown in this Figure and in Fig. 4.

**Fig. S6.**
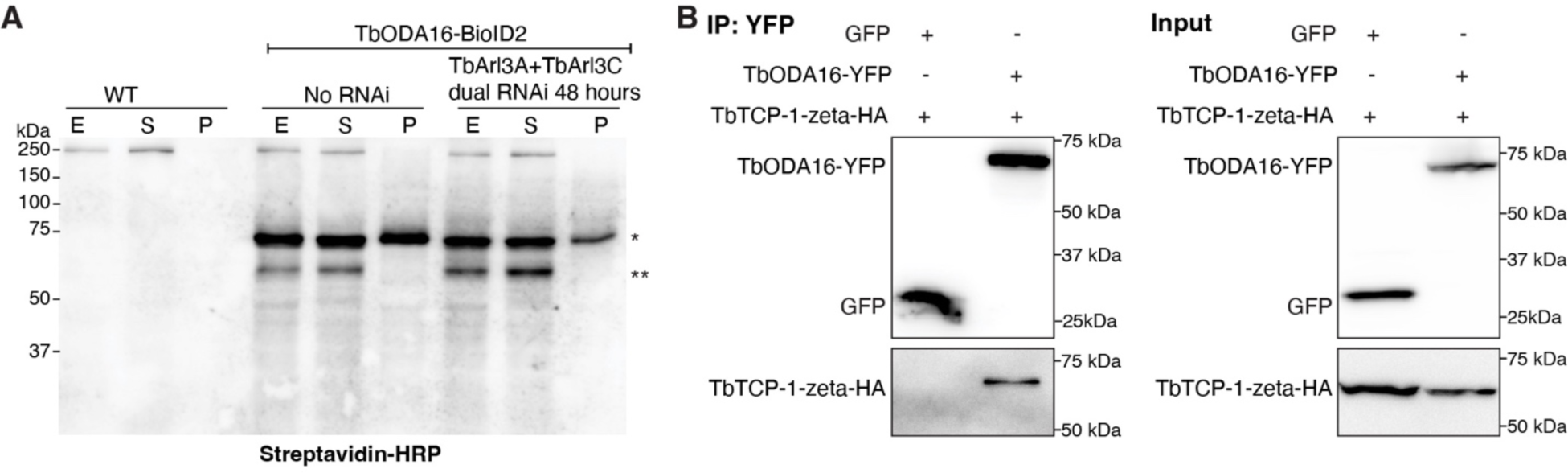
TbODA16 is associated with TRiC. **(A)** Immunoblots probed with streptavidin-HRP revealed the biotinylation profiles of TbODA16-BioID2. The cells were extracted with 1% NP40 in PEM buffer and centrifuged to obtain whole cell (E), detergent soluble (S) and detergent insoluble (P) fractions. WT, wild type cells not expressing TbODA16-BioID2 fusion; no RNAi, cells with expression of TbODA16-BioID2 but not induced for RNAi; TbArl3A+TbArl3C dual RNAi 48 hours, cells with expression of TbODA16-BioID2 and induced for TbArl3A/TbArl3C dual RNAi for 48 hours. * TbODA16-BioID2. ** ∼60 kDa bands corresponding to the size of TRiC subunits. (**B**) Co-IP assays confirming the interaction between TbODA16 and TRiC subunit TCP-1-zeta. Cells expressing GFP were used as a negative control.

## Notes

### Competing Interest Statement

The authors have declared no competing interest.

